# Simple muscle-lever systems are not so simple: The need for dynamic analyses to predict lever mechanics that maximize speed

**DOI:** 10.1101/2020.10.14.339390

**Authors:** A. C. Osgood, G.P. Sutton, S. M. Cox

## Abstract

Here we argue that quasi-static analyses are insufficient to predict the speed of an organism from its skeletal mechanics alone (i.e. lever arm mechanics). Using a musculoskeletal numerical model we specifically demonstrate that 1) a single lever morphology can produce a range of output velocities, and 2) a single output velocity can be produced by a drastically different set of lever morphologies. These two sets of simulations quantitatively demonstrate that it is incorrect to assume a one-to-one relationship between lever arm morphology and organism maximum velocity. We then use a statistical analysis to quantify what parameters are determining output velocity, and find that muscle physiology, geometry, and limb mass are all extremely important. Lastly we argue that the functional output of a simple lever is dependent on the dynamic interaction of two opposing factors: those decreasing velocity at low mechanical advantage (low torque and muscle work) and those decreasing velocity at high mechanical advantage (muscle force-velocity effects). These dynamic effects are not accounted for in static analyses and are inconsistent with a force-velocity tradeoff in lever systems. Therefore, we advocate for a dynamic, integrative approach that takes these factors into account when analyzing changes in skeletal levers.

## Introduction

In this commentary we advocate for an integrative approach to analyzing changes in skeletal lever mechanics. Biomechanists often infer the output speed of an organism’s behaviour from the geometry of its lever system using static or quasi-static analyses which ignore inertial properties of the system (Anderson, 2010; Anderson and Westneat, 2009; Case et al., 2008; Copus and Gibb, 2013; Olivier et al., 2021; Westneat, 1994). In these static conditions, the influence of a simple lever geometry is straightforward. The large mechanical advantage of a crowbar (large input lever and small output lever) for instance, amplifies the force and reduces the velocity of the tip relative to the point of force application. As you decrease the mechanical advantage by applying force closer to the fulcrum, the force amplification (output/input force) decreases while the velocity amplification (output/input velocity) increases. Many have inferred from this that there is a force-velocity tradeoff in lever systems, with greater mechanical advantage (input/output lever arm) producing slower output velocities but greater output forces and smaller mechanical advantage producing faster output velocities but lower output forces (Barel, 1983; M. W. Westneat, 1994). We take issue with this last step of logic: that the ratio of output to input velocity is sufficient to predict the speed of an organism’s movement.

There are two assumptions buried in this inference from static intuitions. First, that one can *increase* the output velocity of a behaviour while *decreasing* the force driving such behaviour. While this is true for a ‘quasi static’ analysis (i.e., an analysis that does not take inertia into account), considering mass results in a significantly more complicated relationship. If one considers mass, and holds all else constant, *decreasing* the distance from the fulcrum at which the force is applied (decreasing mechanical advantage) will *decrease* the torque (Torque=Force*Distance) applied at the joint, *decreasing* output velocity, not increasing it. This is because any increase in output velocity *must* be accompanied by an increase in output force. Thus, for a system with mass, a constant force combined with decreasing mechanical advantage can only decrease the output velocity of the system. Consequently, a mass inclusive analysis and a quasi-static analysis can produce mutually exclusive predictions for the relationship between moment arm and behaviour speed. The quasi-static analysis also makes a second assumption: for the ratio of output to input velocity across a lever to predict the lever speed, there must be a one to one relationship between changes in lever mechanics and changes in output speed (Alfaro et al., 2004). This is not reliably true of lever systems actuated by muscles because this analysis ignores dynamic interactions between muscle force, inertia, and behavioural kinematics (Ackland et al., 2012; Clayton et al., 1998; Dunlop et al., 2004; Galantis et al., 2003; Hoy et al., 1990; Marsh, 1999; McNeill et al., 1972; Nagano and Komura, 2003; Zajac, 1992). These two assumptions are necessary to predict function from lever mechanical advantage alone and, as we shall show, neither of them are valid. If only one message is internalized from this commentary, let it be this: increasing the speed of a system necessitates increasing the force applied, so there cannot be a monatonic trade-off between force and velocity in lever systems.

We are not the first to insist that the performance of biomechanical systems depend on the dynamic interplay between multiple components (Dickinson et al., 2000; Nishikawa et al., 2007) or that the relationship between form and function can be non-linear and complex (Koehl, 1996; Wainwright, 2007). Many have argued against a reductionist approach to understanding biomechanical systems and identified the key roles integrated dynamics play in determining performance. For example, Josephson’s “work loop technique” showed that the same muscle could perform many different functions depending on the temporal pattern of applied strain and activation. Likewise, numerous researchers have recognized that changing lever mechanics can alter the strain rate applied to muscles (Holzman et al., 2008; Richards and Biewener, 2007) or that muscle dynamics alter the function of a lever system (McHenry, 2011; Oufiero et al., 2012; Roberts et al., 2018; Westneat, 2003) or both (Coombs, 1978; Galantis et al., 2003; Marsh, 1999; Zajac, 1992). Yet, while some researchers recognize that a reductionist approach is not sufficient, the dogma of a force-velocity tradeoff in lever systems persists (Arnold et al., 2011; Bergmann and Hare-Drubka, 2015; Brusatte et al., 2012; Patek and Biewener, 2018; Vogel, 2013), particularly in the subfields of fish feeding (Alfaro et al., 2004; Cooper et al., 2017; De Schepper et al., 2008; Evans et al., 2019; James Cooper et al., 2020; McGee et al., 2013; Olivier et al., 2021; Oufiero et al., 2012; Roberts et al., 2018; Turingan et al., 1995; Westneat, 2003) and bird beak biomechanics (Corbin et al., 2015; Herrel et al., 2009). Here we aim to clearly explain why static intuitions that imply a monatonic force-velocity tradeoff are insufficient to predict behaviour speed, and thus an integrative perspective that provides a more subtle understanding of the complex dynamics is necessary.

The need to take an integrative approach will be demonstrated by using dynamic simulations of a simple lever system (via OpenSIM, an open source musculoskeletal modeling program (Seth et al., 2018)) driven by 100% activation of a single muscle. Across 22,572 simulations, we held muscle volume and output lever lengh constant but varied muscle morphology (optimal fiber length, pennation angle and starting normalized fiber length), input lever arm length (thus varying mechanical advantage) and the inertia of the output lever (i.e. resistive forces, See Supplementary Materials for model details). The additional mass was intended to account for the influence of muscle mass, additional body segments or external forces like drag.

By analyzing the results of these simulations we will make 4 arguments for the integrative approach. First, a single lever morphology can produce a wide range of maximum output velocities if muscle properties and resistive forces vary. Second, different lever morphologies can produce identical performance over a wide range of conditions. Third, mechanical advantage is not the most significant determinant of performance in dynamic systems; resistive forces, such as inertia, are more important for determining behaviour speed. Lastly, we look arcross a range of moment arms to provide a mechanistic explanation of how the components of a dynamic lever system interact.

### I: A single lever morphology can produce a wide range of output velocities

To illustrate the range of function possible for a single lever morphology, we subset the results of our simulations to those with a mechanical advantage of 1/8.28. Figure 1 shows the maximum output velocities of 792 simulations with the same mechanical advantage but variable inertia and muscle morphology. The resultant behaviour speeds were as low as 11.15 radians per second and as high as 54.05 radians per second. This implies that the relationship between input and output velocity (moment arm ratio) across a lever system is insufficient to determine the maximum velocity of this system. There is thus not a one to one mapping from lever mechanics to function, as has been suggested (Alfaro et al., 2004). Our simple lever can produce a wide range of output velocities because this is a *lever system* composed of the lever, the driving muscle and the resistive forces. As we will argue throughout this commentary, it is the combination these elements that determine output velocity. This implies that a simple lever system is not, in reality, “simple” and can produce a many to one mapping of morphology to function just as more complex linkages do (Wainwright, 2007).

**Figure 1.**
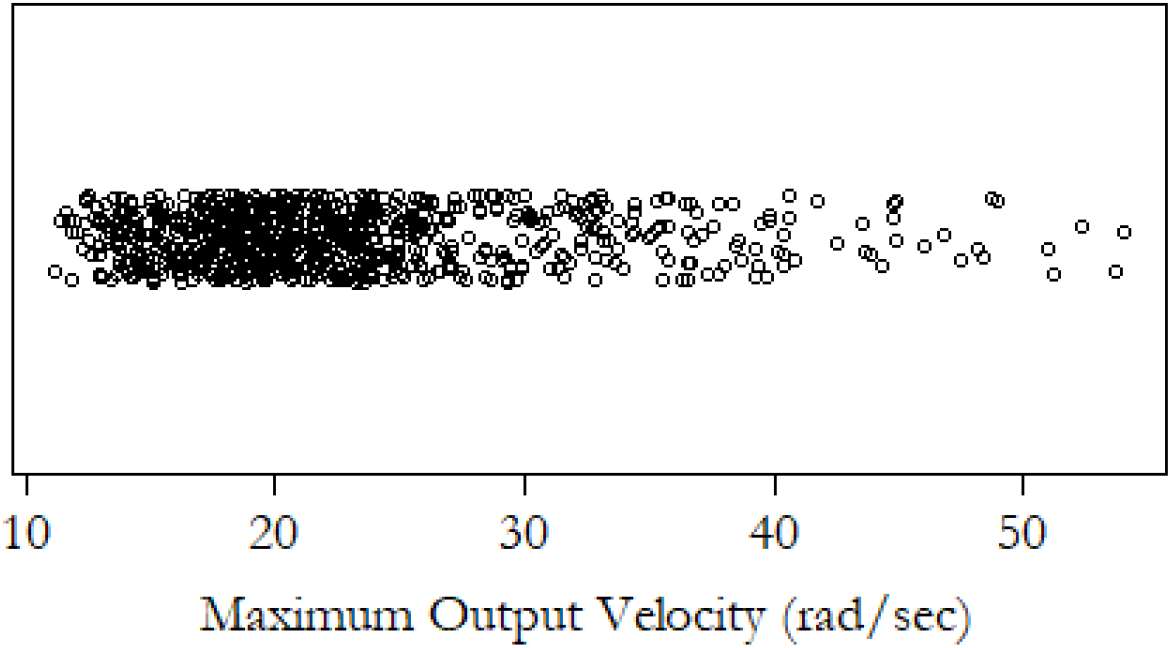
A single lever morphology can produce a wide range of maximum ourput velocities. Here we show the maximum output velocities of 792 unique lever systems with the same mechanical advantage but different muscle (i.e pennation angle, optimal fiber length and starting normalized fiber lengths) and resistive properties. Each dot represents the maximum velocity of a unique lever system. The resistive forces (inertia) acting on the system vary by ±70%, the pennation angle ranges from zero to 40 degrees, and optimal fiber length and normalized muscle start length vary by ±35 and ±20% respectivelyThe data is jittered on the y-axis for clarity.

### II: Drastically different skeletal morphologies can generate the same output kinematics

To contextualize the extent of variability displayed in our first analysis, we compared the possible output kinematics of two drastically different skeletal morphologies with the same variation in muscle and inertial properties. To do so, we subset the results of our simulations to match the mechanical advantage of two example skeletons illustrated in Figure 2A; the forelimb of the horse (mechanical advantage 1:13) and the forelimb of the armadillo (mechanical advantage 1:4 (Smith and Savage, 1955)). While our first analysis aimed to show that a simple lever can produce a one-to-many relationship between form and function, here we aim to illustrate the reverse: that diverse lever morphologies can produce a many-to-one relationship between form and function.

**Figure 2.**
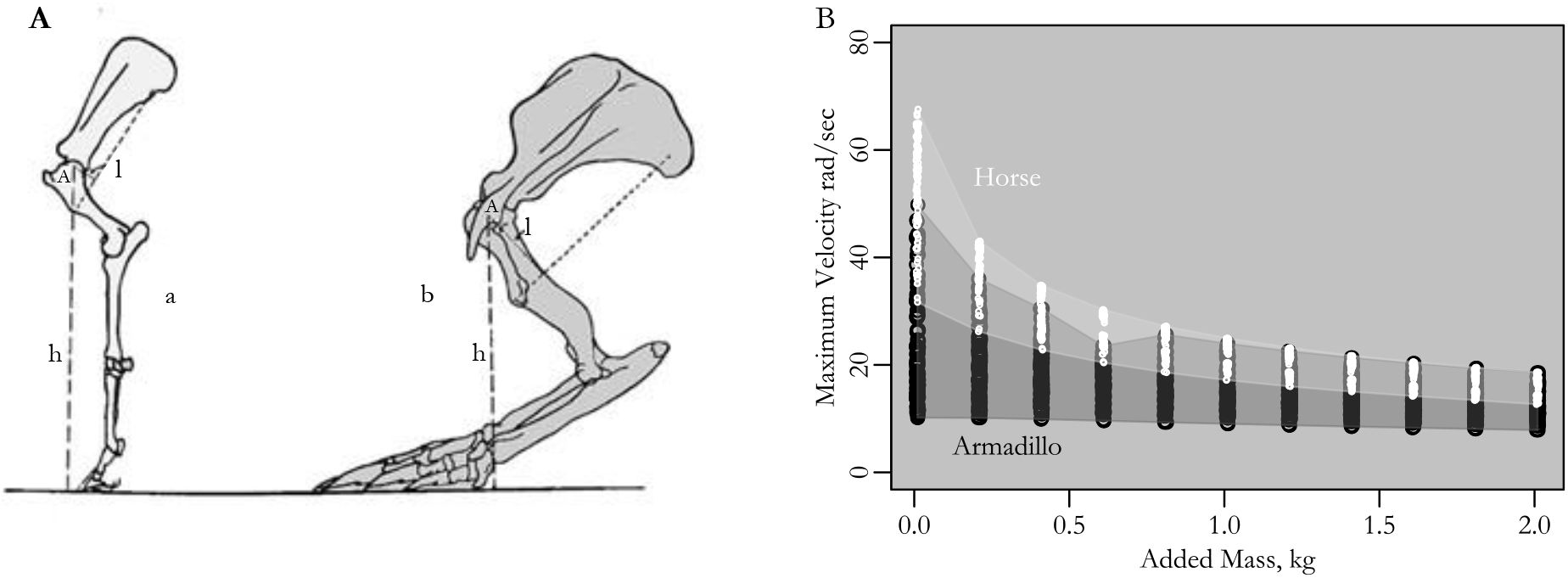
A) Left forelimbs of (a) *Equus* (mechanical advantage 1/13) and (b) *Dasypus* (mechanical advantage ¼), to show the line of action of m. teres major. Text and figure adapted from Smith and Savage 1955. B) The range of output velocities of lever systems with the mechanical advantage of Equus (light grey shaded regions and white points) vs Dasypus (dark grey shaded region with black points) is plotted as a function of the resistive forces acting on the systems. The functional overlap for the two different lever morphologies is broad and increases with increasing resistance.

In Figure 2B, we plot the possible output velocities of the two lever systems described above against the resistance force encountered. Three main points should be taken from this figure. The first reinforces the conclusions of our last analysis, namely that the lever mechanics of the horse and armadillo can produce a wide range of output velocities, both across all conditions and even when encountering the same resistive forces (described as ‘Added Mass’ in Figure 2B). Secondly, for every driven mass, there is an overlapping region of output performance where the two skeletal morphologies produce the same velocity. Changes in mechanical advantage alone do not definitively determine the function of the system. Since variations in output velocity for any driven mass are the result of changes in muscle morphology alone, the overlapping regions are only possible because lever mechanics can be offset by changes in muscle morphology (Lee and Piazza, 2009; Zajac, 1992). This is a viable biological path as skeletal and muscle morphologies have been shown to evolve distinctly (Roberts et al., 2018).

The last point to be taken from Figure 2B is that as resistive forces (inertia) decrease, the region of overlapping output between the two lever systems decreases. This means that changes in mechanical advantage have a greater influence on the function of a system when the resistive forces are small. Thus, the extent of muscle variation needed to compensate for changes in mechanical advantage will be very high for small resistive forces and decrease for larger masses. Taking this trend to its logical extreme, when resistive forces are ignored (as in static and quasi-static analyses), the maximum velocity of a system would appear highly sensitive to changes in mechanical advantage. Ignoring resistive forces, however, is not a reasonable simplification to make given their impact on maximum velocity. Therefore, the one-to-one mapping between changes in lever mechanics and function assumed in quasi-static analyses does not capture the more complex dynamics of lever systems. Integrative analyses of variation in skeletal morphology may be necessary to avoid significant errors when studying the morphological variation that enables animals to move quickly.

### III: Mechanical advantage isn’t the most important factor in determining output velocity

Our first two analyses suggest that other factors, such as the resistive forces, may be more important than mechanical advantage in determining the performance of a lever system. In this third analysis, we quantify the relative contribution of muscle properties, mechanical advantage, and resistive forces to the maximum velocity of our lever system. We first built linear statistical models for each of the morphological elements in our lever system, (i.e. muscle force capacity, muscle pennation angle, starting fiber length, mechanical advantage, and added mass (i.e. inertia)) and evaluated the explanatory power of each predictor individually (R Core Team, 2017).

As we have repeatedly argued for the need to take an integrative approach to analyzing these systems, we also built a multivariable linear regression model including all of the parameters described above and their interaction effects. To determine which subset of the full list of morphological parameters has the most explanatory power we performed a stepwise AIC (Akaike Information Criterion) model comparison. We hoped first, to quantify the most significant contributors to performance when the system is taken as a whole, then to compare the explanatory power of this full model to the best individual predictor. See Supplemental Materials for additional model details and results.

When comparing the explanatory power of individual predictors, the adjusted R^2^ values in our first analysis reveal output velocities to be the most sensitive to resistive force (i.e. inertia). Resistive forces explain 45% of the variation in the maximum velocity of our systems while mechanical advantage, though the second most significant predictor, only explains 5%. The relative importance of mass and moment arm can be seen visually in Figure 3A where we compare the variation in maximum velocity across different moment arms for two different resistive forces driven by the same muscle. The figure illustrates three points. First, in conjunction with the statistical results, Figure 3A suggests that changing limb inertia can have drastically larger effects on system kinematics than changing mechanical advantage. This implies that the major error in quasi-static analyses is that they do not include the aspect of the system (i.e. inertia) that most substantially alters kinematics.

**Figure 3.**
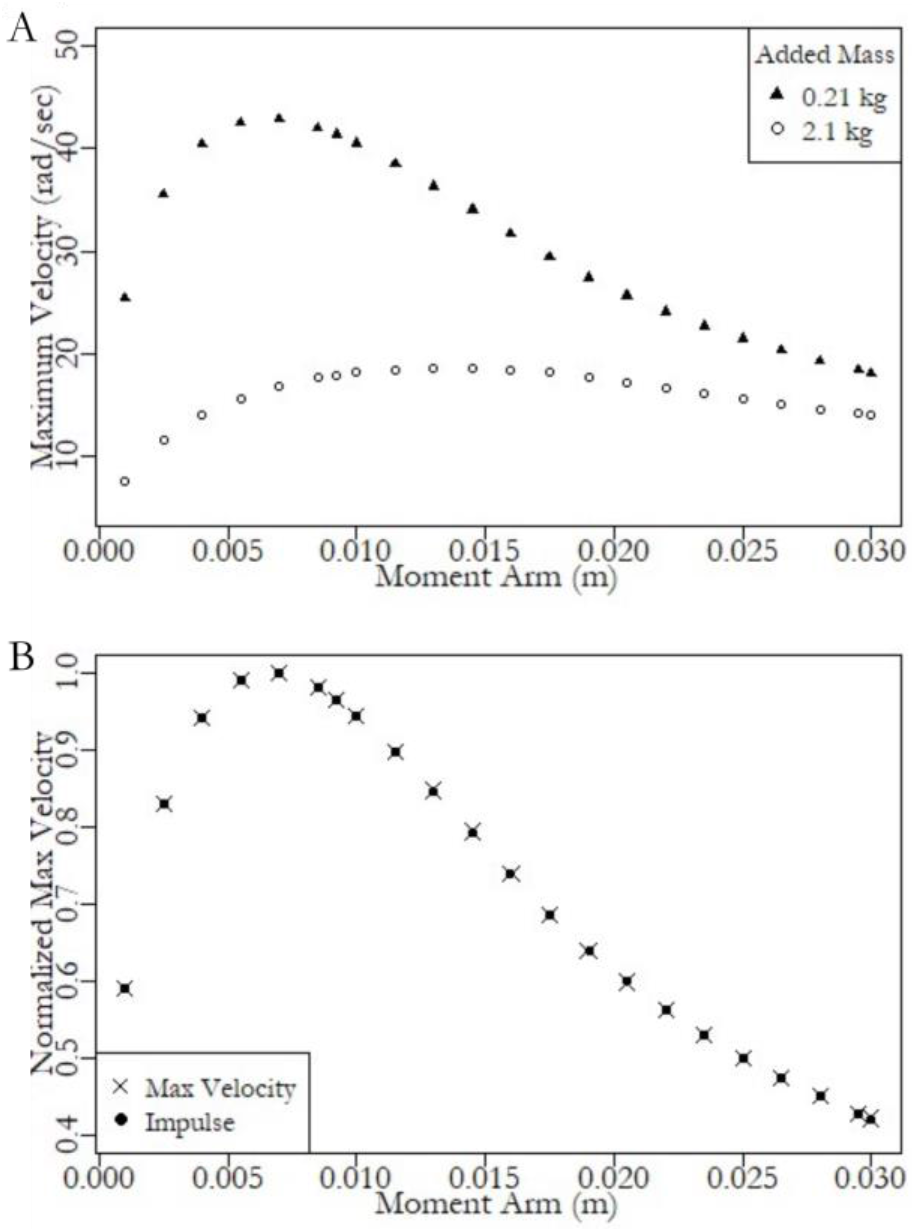
A) The maximum output velocities for lever systems driven by the same muscle but across a range of moment arms and encountering different resistive forces (black circles: lever inertia + 0.01 kg added mass, grey circles: lever inertia + 2 kg added mass). Note that output velocity does not increase linearly with decreasing mechanical advantage. Rather, there is an optimum mechanical advantage and that optimum value changes with resistive forces. Thus, the same change in mechanical advantage can either increase or decrease output velocity in different conditions. Further, the change in output velocity is more sensitive to changes in resistive forces than changes in moment arm in many conditions. B) Total output impulse and maximum system velocity exactly coreespond when normalized by their maximum values. Any increase in output velocity must be accompanied by an increase in output force.

Second, notice the inverted U-shape of the curves in Figure 3A which would not be predicted from quasi-static analyses. Figure 3A makes it particularly clear that changes in output velocity do not change *monotonically* with mechanical advantage. Specifically, depending on the muscle and inertial properties, the same change in mechanical advantage could *increase* or *decrease* the maximum output velocity. Importantly, for systems with mass, there is often an optimal mechanical advantage that will maximize output speed for a given set of muscle and inertial conditions (Coombs, 1978). Lastly, in agreement with the results of our previous analysis, the magnitude of the functional change resulting from an adjustment in lever mechanics varies as a function of both mechanical advantage and resistive forces.

As expected from the integrated nature of these systems, the full model, incorporating all of the morphological parameters, can explain 69% of the variation in maximum velocity. To reiterate, the best individual model (taking only mass into account) only explained 44% of the variation. This again highlights the integrated nature of these systems and the need to study them as a whole.

### IV: Decreasing moment arm does not necessarily increase output velocity

We have tried to show the potential problems with using static or quasi-static analyses to predict function from just skeletal morphology. But pointing out problems without providing alternatives is not sufficient. A better goal, one we attempt, is to offer an alternative framework to drive intuitions about the influence of changes in dynamic lever systems that can provide a more subtle understanding of how components work together dynamically. In this section, we present a more detailed results to illustrate the dynamic interactions that influence performance metrics as lever mechanics vary. Specifically, in Table 2, we show the results of simulations with the smallest, largest and optimal (i.e. producing highest output velocity, see figure 3A) for a single driven mass and muscle morphology.

**Table 1:** Statistical models comparing predictive power of individual morhological variables (shaded grey region) with a multivariate model including interaction effects.

| Model | coeff | t-value | Adj. R<sup>2</sup> | P-value | AIC | delAIC |
| --- | --- | --- | --- | --- | --- | --- |
| **Null** | 21.83 |  |  | <2e-16 | 169644 | 26651 |
| **Pennation Angle_rad** | 5.69 | 20.87 | 0.0189 | <2.2e-16 | 169215 | 26221 |
| **Optimal Fiber Length** | 0.75 | 18.60 | 0.01505 | <2.2e-16 | 169,303 | 26,309 |
| **Starting n. Fiber length** | 9.89 | 23.70 | 0.02425 | <2.2e-16 | 169,091 | 26097 |
| **Moment arm_mm** | -0.27 | -35.8 | 0.05368 | <2.2e-16 | 168,400 | 25406 |
| **Added Mass_kg** | -10.96 | -135.0 | 0.4469 | <2.2e-16 | 156,278 | 13284 |
| **MomentArm \* Added Mass\* Pennation Angle \* n.Fiber Length \* Optimal Fiber Length** |  |  | 0.6933 | <2.2e-16 | 142,994 | 0 |

**Table 2:** Morphological parameters and output from simulations varying moment arm but holding all else constant.

| MA (mm) | Varying moment arm |  |  |
| --- | --- | --- | --- |
|  | Min | Opt | max |
|  | 1.0 | 7.0 | 29.5 |
| **Added Mass (kg)** | 2.01 | 2.01 | 2.01 |
| **Starting Normalized Fiber Length** | 1 | 1 | 1 |
| **Optimal Fiber Length (m)** | 0.07 | 0.07 | 0.07 |
| **Pennation Angle (deg)** | 0 | 0 | 0 |
| **Maxium Output Velocity (Rad/s)** | 10.00 | 17.60 | 9.52 |
| **Max Muscle Velocity (m/sec)** | 0.20 | 2.46 | 5.62 |
| **Muscle contraction distance/Optimal Fiber Length** | 0.05 | 0.37 | 0.56 |
| **In/Out Velocity Ratio** | 1.00e-03 | 7.00e-03 | 0.29e-03 |
| **Muscle Impulse (Ns)** | 1.17e+07 | 2.94e+06 | 3.78e+05 |
| **Output Impulse (Ns)** | 9.60e+04 | 1.69e+05 | 9.14e+04 |
| **Total muscle work (Nm)** | 3.06e+04 | 5.39e+04 | 1.05e+04 |
| **Total Output Power N/s** | 5.86e+04 | 1.81e+05 | 5.30e+04 |
| **Muscle Power/ Output Power** | 1 | 1 | 1 |
| **Time to Maximum Velocity (s)** | 0.53 | 0.26 | 0.31 |

A static analysis would predict that decreasing moment arms would *increase* output velocities. Instead, we found decreasing moment arm can sometimes *decrease* output velocity, as illustrated in Figures 3A&B. For instance, when comparing the maximum velocities from Table 2, decreasing the moment arm from 7 to 1.1 mm drops maximum velocity from 17 to 10 rad/s. To make sense of this, it is important to consider how lever arms affect torque. As stated earlier, in worlds where objects have mass, one cannot increase the output velocity of a limb without increasing the force driving it as illustrated in Figure 3B. Any increase in velocity must be driven by an increase in torque. Holding all else constant, a decrease in moment arm decreases the torque available to drive the motion. Thus, in a dynamic system driven by an ideal actuator (one that could produce constant force at any speed), a decrement in moment arm would decrease both output velocity and output force. A second factor also limits output velocity at small moment arms. In a system with a constrained range of motion, as we have here, a small mechanical advantage also limits the work a muscle can produce. As moment arms decrease, the available contractile distance over which a muscle can apply force will also decrease. Again, for an ideal actuator, this decreases the energy (force*distance) that can be applied, and thus decreases maximum output velocity. Output velocity, then, drops off as force is applied closer to the point of rotation primarily because torques and muscle displacements are both limited by a small input lever arm. Consequently, the intuition that low mechanical advantage necessarily improves output velocity is often completely backwards.

The interesting question should really be, then, why does output velocity also drop at larger moment arms despite increasing mechanical advantage? In our models, this drop off is primarily due to muscle-force velocity effects, in agreement with predictions by others (Ilton et al., 2018; Lee and Piazza, 2009; Nagano and Komura, 2003; Sutton et al., 2019; Zajac, 1992). In Table 2, this can best be seen by noticing the increase in maximum muscle velocity as muscle impulse decreases with increasing moment arm. Levers effectively alter the resistive forces and thus the strain pattern acting on the muscle. Thus, we can explain the shape of the output velocity-moment arm plot as the interaction of two opposing factors: those decreasing velocity at low mechanical advantage (low torque and muscle work) and those decreasing velocity at high mechanical advantage (force-velocity effects). These two effects will cause each lever system to have an optimum mechanical advantage that minimizes these opposing factors, as noted by others (Galantis et al., 2003; Josephson, 1985; Zajac, 1992).

It is important to note that the ratio of input to output velocity still changes linearly with mechanical advantage (See Table 2). Yet, as we suggest in the introduction, this ratio does not predict the maximum velocity of the whole motion. The output velocity is, instead, maximized at an intermediate mechanical advantage value (See Figures 3A&B).

#### What could replace quasi-static analyses?

Our main goal with this commentary is to change the conceptual framework used to think about the relationship between form and function in lever systems. Specifically, we argue for a need to move away from the intuitions derived from reductionist static analyses which imply a force-velocity tradeoff in lever mechanics. We propose shifting our thinking to an integrative framework that acknowledges the dynamic interactions between the muscle, lever mechanics and resistive forces (i.e. inertia) acting on the system. Within this new framework, there is a tradeoff between constraints that limit output speed by limiting torque at low mechanical advantages and constraints that limit output speed by limiting muscle force at high mechanical advantage through muscle force-velocity properties.

In addition to changing the way we think about these systems, we also hope to provide an example of how best to improve the accuracy of predicting the functional conseques of variations in skeletal morphology. Given the highly integrated nature of these systems, we recommend performing dynamic analyses that include direct measurements of muscle properties (pennation angle, starting fiber length, ofl) and resistive forces to most accurately predict the influence of changes in lever mechanics on kinematics. There are several open source musculoskeletal modeling programs available to do this (Seth et al., 2018; Todorov et al., 2012) and published examples of in house built models or studies using opensource software abound (De Schepper et al., 2008; Farris et al., 2014; Hutchinson et al., 2015; Ilton et al., 2018; Richards and Eberhard, 2020; Roberts, 2003). See the supplemental materials for example code used in this manuscript.

If one does not have access to extant specimens from which to measure muscle and inertial properties, we suggest building musculoskeletal-models that match the observable lever mechanics and performing sensitivity analyses (Ackland et al., 2012; Anderson et al., 2007; Hutchinson, 2004). This can be done by varying unknown muscle and inertial properties through monte-carlo simulations (similar to our first two analyses) to get a measure of the uncertainty in estimates of function. With this approach, one could test hypotheses about the functional consequences of changes in skeletal morphology while accurately capturing the uncertainties. Lastly, for back of the envelope estimates, our results suggest that shifting the focus from changes in lever mechanics to changes in inertial properties and external resistive forces would result in more accurate predictions of system maximum speed.

In summary, we suggest that it is not appropriate to assume a force-velocity tradeoff in muscle-driven lever systems when looking across a whole motion because increasing output velocity always requires increasing output force. We argue that inferences from changes in lever mechanics alone to changes in function are error prone. This is for two reasons. First, quasi-static analyses do not incorporate the most sensitive parameter, namely inertial effects. Second, the influence of variation in individual parameters are highly interdependent. Thus, analyses that predict changes in output from changes in a single input parameter will often predict inaccurate functional consequences. Quasi-static analyses are, however, correct for systems in which the inertial effects are very small because acceleration is negligible (for example, bite forces in fish). As inertial effects get larger, however, the predictive power of quasi-static analyses weakens.

The analysis of lever systems which include the dynamic muscle and inertial effects yield an improved quantitative framework to evaluate form and function. We hope this commentary will both provide warning of the possible range of errors from the current framework and a roadmap for how best to generate dynamic analyses to overcome these limitations.

## Supporting information

Supplemental Methods Descriptions

## Acknowledgements

We’d like to thank Manny Azizi and Jeff Olberding for reading and commenting on an early draft of this manuscipt.

## Data Availability

The code used in the project and the generated data are available at DataDryad https://datadryad.org/stash/share/AqKLIaK7A1NgxKoLYGD7SX7Nab92qANnrhrInx-9RRc.

