## Supplemental Methods Descriptions for "Simple muscle-lever systems are not so simple: The need for dynamic analyses to predict lever mechanics that maximize speed"

**Supplemental Materials**

1. **Musculoskeltal model details**

To quantify the relationship between force velocity properties, mechanical advantage, inertia, and maximum output velocity in muscle driven lever systems, we built a computational model of a simple lever system using OpenSim V. 4.0[15], scripting in MATLAB (MathWorks; Natick, MA). Our model was composed of an output lever (length 0.12 m, width and height 0.04 m) that pivoted around a pin joint. A stand connected the pin-joint to the world and a muscle originated on the stand, wrapping around a wrap cylinder beforeattaching to the lever (Figure A1). These dimensions were chosen to scale the model to approximate the tarsometatarsus of the guinea fowl since detailed experimental data was available(Cox et al., 2019). The diameter of the wrapping surface defined the input lever (moment arm) of the muscle acting at this joint and allowed the muscle to change length and rotate the lever without altering the line of action. The muscle implemented was a Millard2012EquilibriumMuscle muscle model using default length-tension and force-velocity (Millard et al., 2013) values and a non-compliant tendon. The lever had the inertial properties of a rectangular rod defined by its volume (0.02 x 0.02 x 0.12 m) with a density of water (997 kg m^-3^). Gravity was not included in the model to isolate the effects of resistive forces without introducing variations due to changing effective mechanical advantage(Roberts, 2003). To simulate driving a mass, we welded a mass to the end of the lever in the shape of a sphere with the density of water. We constrained the lever’s range of motion to 150 degrees with a joint limiting force stiffness of 20, damping of 5 and transition range of 2.5 degrees.

We generated 22572 modifications of this model with different muscle optimal fiber lengths (OFL:(0.05:0.01:0.1 m), moment arms (0.0005:0.00125:0.02 and 0.12/13 m), starting normalized muscle lengths (0.8,1.1,1.4), pennation angles (0:10:40) and driven masses (0.01: 0.02: 2 kg). Our range of added masses span those an animal might encounter during the swing phase (only driving the limb inertia) and stance phase (accelerating the center of mass of the body) of locomotion.

To compensate for the change in muscle-tendon unit length as moment arm changed, we adjusted the tendon slack length for each model such that the equilibrium normalized fiber length for the passive muscle was within 0.01 units of a fixed starting fiber length. We held the volume of the muscle constant across changes in optimal fiber length. As the maximum isometric force of a muscle is a function of its cross-sectional area, an iso-volumetric muscle implies that the force capacity, *F_max_*, is inversely related to optimal fiber length, OFL, such that

$F_{max}OFL = \frac{M T_{s}}{\rho}$,

where *M* is muscle mass, T_s_ is the specific tension of the muscle and $\rho$ is muscle density. The muscle was modeled on the sum of the lateral and medial gastrocnemius muscles of the guinea fowl[16], with a muscle mass of 17g, a specific tension of 3e5 N/m^2^ and a muscle density of 1060 kg/m^3^. For each modified model, we simulated motion resulting from 100% activation of the muscle model across the full range of motion of the joint in 0.0005 second time steps.

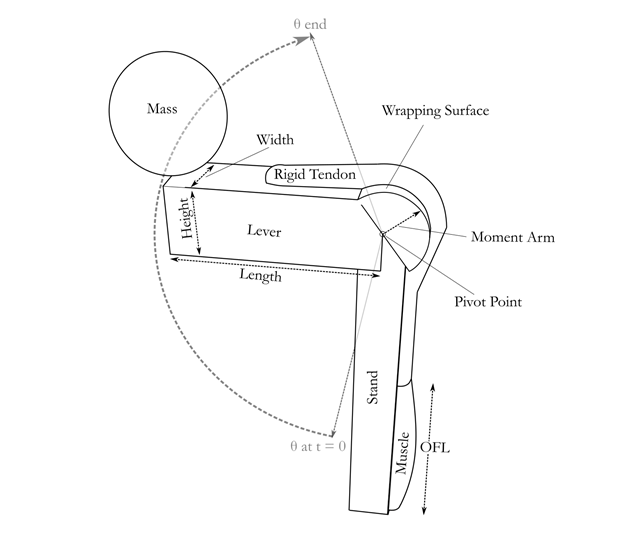

Figure A1. Schematic of the simple lever system built in Opensim to model the dynamic interactions between muscle properties, lever mechanics and resistive forces. The contraction of the muscle applies a torque at the joint a distance from the pivot point defined by the diameter of the surface that the muscle wraps around.

At each time step we extracted the joint angle, angular velocity and acceleration of the lever as well as the muscle active-fiber force, fiber length, and fiber velocity. We trimmed this data to the time range over which the velocity of the mass was increasing. To distinguish the torque required to accelerate the lever from that available to accelerate the mass, we calculated them separately. The force acting to accelerate the mass was calculated from the torque accelerating the mass,$\tau_{m}$, and the distance from the center of the mass to the pivot, *d*.

$$F_{m}=\frac{\tau_{m}}{d}$$

The torque acting to accelerate the mass was calculated from the acceleration of the mass and its moment of inertia rotating around the pivot, using the parallel axis theorem;

$$\tau_{m} = \frac{2}{5} mr^{2}+md^{2}$$

where d is the distance from the center of mass to the pivot point, r is the diameter of the added mass, and *m* is the mass of the added mass. The torque acting to accelerate the lever was similarly calculated.

Additionally, impulse of the muscle and impulse applied to the mass were calculated as the integral of the active fiber force of the muscle and the force applied to the mass, respectively.

2. **Statiscal Model Details**

We evaluated the predictive power of morphological variation on the maximum velocity of our simulations with a multivariate linear model with muscle pennation angle, optimal fiber length, normalized start length, added mass and moment arm as predictor variables including interaction effects.

fullModel = lm(maxVel~momentArm_mm*pennationAngle*mass*ofl*initFiberLength,data=es)

After a stepwise AIC forewards and backwards model comparison (stepAIC, (Venables and Ripley, 2002), the best full model included the following parameters and interactions:

Full Model: lm(formula = maxVel ~ momentArm_mm + pennationAngle + mass +

ofl + initFiberLength + momentArm_mm:pennationAngle + momentArm_mm:mass +

pennationAngle:mass + momentArm_mm:ofl + pennationAngle:ofl +

mass:ofl + momentArm_mm:initFiberLength + pennationAngle:initFiberLength +

mass:initFiberLength + ofl:initFiberLength + momentArm_mm:pennationAngle:mass +

momentArm_mm:pennationAngle:ofl + momentArm_mm:mass:ofl +

pennationAngle:mass:ofl + momentArm_mm:pennationAngle:initFiberLength +

momentArm_mm:mass:initFiberLength + pennationAngle:mass:initFiberLength +

momentArm_mm:ofl:initFiberLength + pennationAngle:ofl:initFiberLength +

mass:ofl:initFiberLength + momentArm_mm:pennationAngle:mass:ofl +

momentArm_mm:pennationAngle:ofl:initFiberLength + momentArm_mm:mass:ofl:initFiberLength,

data = es)

Residuals:

Min	1Q	Median	3Q	Max

-23.9431 -2.4694 -0.4932	2.0247 28.7966

Coefficients:

Estimate Std. Error t value Pr(>|t|)

(Intercept)	20.180918	5.219030	3.867 0.000111 ***

momentArm_mm	-0.890710	0.288057 -3.092 0.001990 **

pennationAngle	33.275829	9.028900	3.685 0.000229 ***

mass	-12.086132	3.551568 -3.403 0.000668 ***

ofl	1.387833	0.674952	2.056 0.039775 *

initFiberLength	19.301599	5.088855	3.793 0.000149 ***

momentArm_mm:pennationAngle	-0.332156	0.494229 -0.672 0.501546

momentArm_mm:mass	0.691289	0.194818	3.548 0.000388 ***

pennationAngle:mass	-18.018224	2.648200 -6.804 1.04e-11 ***

momentArm_mm:ofl	-0.001766	0.037447 -0.047 0.962381

pennationAngle:ofl	-3.527573	1.157873 -3.047 0.002317 **

mass:ofl	-0.035628	0.456554 -0.078 0.937800

momentArm_mm:initFiberLength	-0.844969	0.280812 -3.009 0.002624 **

pennationAngle:initFiberLength	-18.872733	8.640818 -2.184 0.028962 *

mass:initFiberLength	0.773192	3.416199	0.226 0.820946

ofl:initFiberLength	-0.545399	0.657958 -0.829 0.407155

momentArm_mm:pennationAngle:mass	-0.071427	0.122019 -0.585 0.558298

momentArm_mm:pennationAngle:ofl	0.178484	0.064261	2.777 0.005483 **

momentArm_mm:mass:ofl	-0.045115	0.025327 -1.781 0.074878 .

pennationAngle:mass:ofl	1.185313	0.285811	4.147 3.38e-05 ***

momentArm_mm:pennationAngle:initFiberLength 0.900117	0.472388	1.905 0.056733 .

momentArm_mm:mass:initFiberLength	0.007829	0.187230	0.042 0.966645

pennationAngle:mass:initFiberLength	7.839379	1.475662	5.312 1.09e-07 ***

momentArm_mm:ofl:initFiberLength	0.080957	0.036505	2.218 0.026586 *

pennationAngle:ofl:initFiberLength	1.396374	1.106705	1.262 0.207055

mass:ofl:initFiberLength	-0.925651	0.438772 -2.110 0.034900 *

momentArm_mm:pennationAngle:mass:ofl	-0.047335	0.015863 -2.984 0.002848 **

momentArm_mm:pennationAngle:ofl:initFiberLength -0.123628 0.061419 -2.013 0.044142 *

momentArm_mm:mass:ofl:initFiberLength	0.040812	0.024341	1.677 0.093617 .

---

Signif. codes: 0 ‘***’ 0.001 ‘**’ 0.01 ‘*’ 0.05 ‘.’ 0.1 ‘ ’ 1

Residual standard error: 5.742 on 22543 degrees of freedom

Multiple R-squared: 0.6937,	Adjusted R-squared: 0.6933

F-statistic: 1823 on 28 and 22543 DF, p-value: < 2.2e-16
